# The ERK5 MAP kinase regulates annexin complexes and membrane dynamics in embryonic stem cells

**DOI:** 10.1101/2024.07.03.598211

**Authors:** Helen A. Brown, Ludivine Guillet, Charles A. C. Williams, Hayley Shaw, Houjiang Zhou, Diana Rios-Szwed, Rosalia Fernandez-Alonso, Liam McMulkin, Marios P. Stavridis, Greg M. Findlay

## Abstract

The ERK5 MAP kinase signalling pathway controls key processes in mouse Embryonic Stem Cells (mESCs) maintenance, including expression of naïve pluripotency genes and a transcriptional programme for stem cell rejuvenation. However, quantitative proteomics suggests that ERK5 signalling controls levels of proteins implicated in diverse biological processes. Here, we show that Annexin A2 (ANXA2) and its accessory protein S100A10, which are implicated in regulation of membrane dynamics, are major targets of the ERK5 signalling pathway in mESCs. ERK5 activation promotes expression of the ANXA2/S100A10 protein complex via transcriptional induction of the KLF2 pluripotency factor. Furthermore, either ERK5 signalling or ectopic expression of the ANXA2/S100A10 complex promotes formation of membrane protrusions in mESCs. Our data therefore identify a new function for the ERK5 pathway in regulation of an Annexin complex and membrane dynamics, which may have implications for development and disease.

## Introduction

Embryonic Stem Cells (ESCs) have the capacity to self-renew or differentiate into any cell type in the adult body, a property known as pluripotency (Nichols & Smith, 2009). The fundamental regulatory mechanisms which govern pluripotency are therefore of intense interest to exploit pluripotent cells in regenerative therapeutics. Cellular signalling networks play a critical role in ESC fate decisions, implementing specific gene expression signatures to define ESC developmental choice (Fernandez-Alonso *et al*, 2017a). Although many of the key signalling pathways that control ESC decision-making have been identified, e.g. (Smith *et al*, 1988; Ying *et al*, 2003; Ying *et al*, 2008), it remains a challenge to elucidate their molecular functions in ESCs.

The ERK5 MAP kinase signalling pathway was identified as a key regulator of early embryonic gene expression (Brown *et al*, 2021), transition between naïve and primed pluripotency in mESCs (Williams *et al*, 2016) and somatic cell reprogramming to pluripotency (Morikawa *et al*, 2016). ERK5 uniquely encodes a kinase and putative transcriptional activation domain (Kasler *et al*, 2000) and promotes the naïve state by driving expression of pluripotency genes (Williams *et al*., 2016). However, efforts to identify wider molecular targets and biological functions of the ERK5 pathway in mESCs are in their infancy. In this regard, quantitative proteomics indicates that ERK5 signalling regulates expression of a highly specific cohort of proteins associated with diverse biology (Brown *et al*., 2021). Indeed, the ERK5-KLF2 signalling axis regulates expression of ZSCAN4 (Brown *et al*., 2021), which plays a key role in regulating expression of early embryonic genes, attainment of naïve pluripotency and stem cell ‘rejuvenation’. These findings provide the first molecular insight into ERK5 functions in mESC proteome patterning. However, the biological functions of other ERK5-dependent targets within the mESC proteome have yet to be investigated.

Here, we show that the Annexin accessory protein S100A10, which is implicated in regulation of cytoskeletal and membrane dynamics, is a major target of the ERK5 signalling pathway in mESCs. ERK5 activation promotes expression of the ANXA2/S100A10 protein complex via transcriptional induction of the KLF2 pluripotency factor. Furthermore, either activation of ERK5 signalling, or mimicking the effect of ERK5 signalling on ANXA2/S100A10 by overexpression, promotes formation of membrane protrusions in mESCs. Our data therefore identify a new function for the ERK5 pathway in regulation of an Annexin complex and mESC membrane dynamics, which has implications for understanding the role of ERK5 in development and diseases such as cancer.

## Results

### Quantitative proteomic profiling identifies S100A10 and ANXA2 as major targets of ERK5 signalling in mouse embryonic stem cells (mESCs)

Previously, we developed a quantitative proteomics workflow to identify proteins whose expression is regulated by ERK5 signalling in mESCs, which uncovered a function for ERK5 in induction of the stem cell rejuvenation factor ZSCAN4 and other early embryonic genes (Brown *et al*., 2021). However, this experimental approach identified and quantified a total of 8732 proteins, of which ERK5 signalling acutely regulates a selective cohort of 56 proteins with diverse biological functions (Fig. 1A). Amongst the top hits identified in this screen, S100A10 protein abundance increases >2 fold upon ERK5 activation by expression of a constitutively-active MEK5 mutant (MEK5DD), and this is partially reversed by treatment with the selective ERK5 inhibitor AX15836 (Fig. 1B).

**Figure 1:**
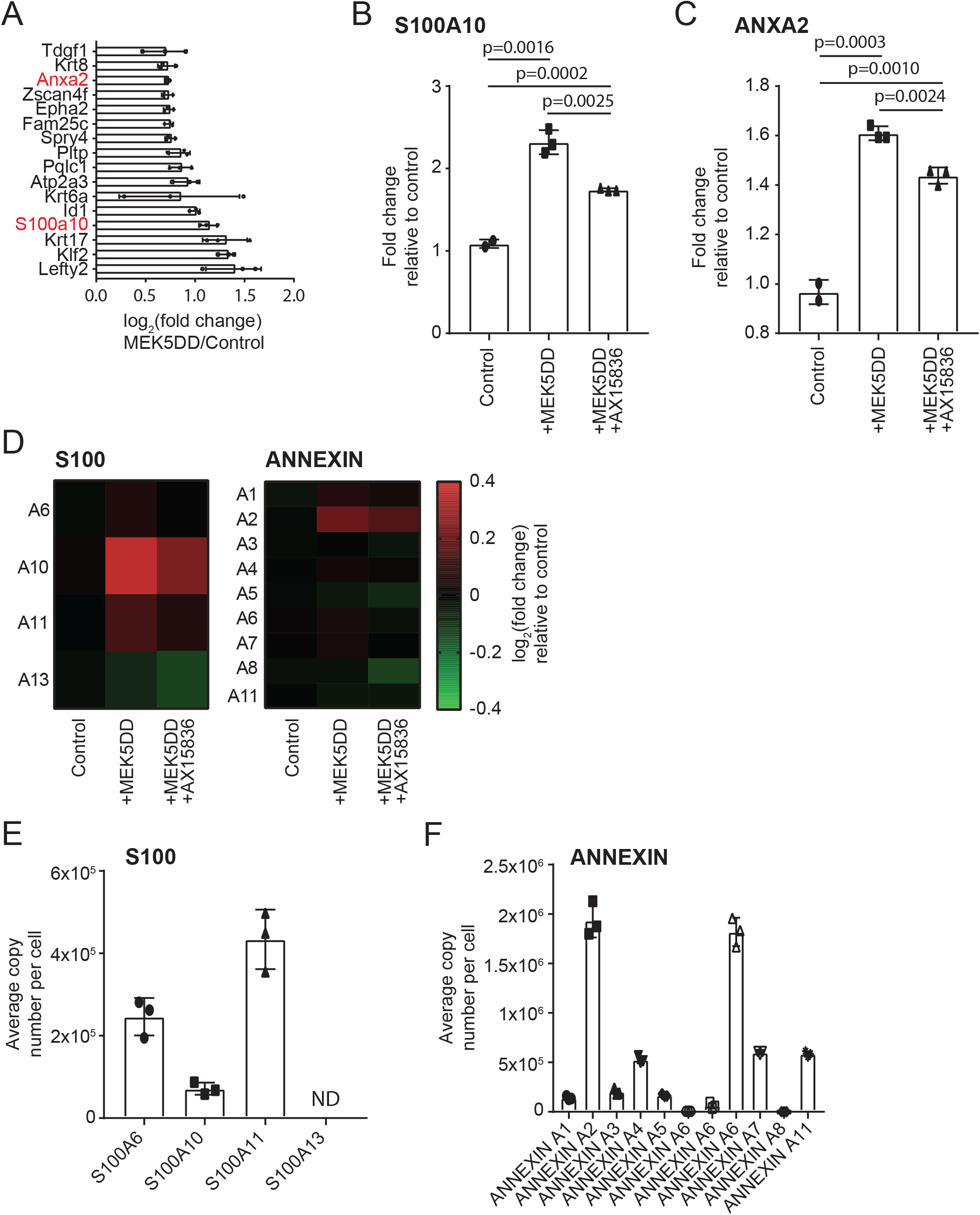
S100A10 and ANXA2 gene expression is specifically regulated by ERK5 signalling in mouse embryonic stem cells (mESCs). (A) Total proteome mass spectrometry determination of log_2_ (fold change) in protein abundance comparing control mESCs to ERK5 pathway activation (MEK5DD) for the proteins that exhibit the greatest fold change. Each data point represents a biological replicate compared to the mean of control conditions. Original dataset was reported in (Brown *et al*., 2021). Error bars indicate standard deviation (n=3). (B) Total proteome mass spectrometry determination of log_2_ (fold change) in S100A10 protein abundance comparing control mESCs to ERK5 pathway activation (MEK5DD) and ERK5 pathway activation followed by ERK5 inhibition (+MEK5DD+AX15836). ERK5 inhibition by AX15836 was for 24 h at a final concentration of 10 μM. Original dataset was reported in (Brown *et al*., 2021). Error bars indicate standard deviation (n=3). (D) Heatmap displaying fold change in S100 and Annexin family protein abundances in mESCs detected and quantified by total proteome mass spectrometry. Original dataset was reported in (Brown *et al*., 2021). S100A10 (left) and ANXA2 (right) protein abundance in control samples was set to 1.0 and relative abundances calculated for control, ERK5 pathway activation (+MEK5DD) and ERK5 pathway activation followed by ERK5 inhibition (+MEK5DD +AX15836). ERK5 inhibition by AX15836 was for 24 h at a final concentration of 10 μM. (E) Average copy number per cell of detected members of the S100 protein family in mESCs was determined using total proteome mass-spectrometry (Fernandez-Alonso *et al*, 2017b) and the histone ruler method (Wisniewski *et al*., 2014). ND = not determined due to incomplete data. (F) Average copy number per cell of detected members of the Annexin protein family in mESCs was determined using total proteome mass-spectrometry (Fernandez-Alonso *et al*., 2017b) and the histone ruler method (Wisniewski *et al*., 2014). Multiple ANXA6 proteoforms were identified and quantified in this dataset.

S100A10 is a member of the S100 family of EF-hand calcium binding proteins, which form heterotetrameric complexes with ANXA2 (also known as Annexin A2/II) (Rety *et al*, 1999; Rintala-Dempsey *et al*, 2008) and function in regulation of membrane dynamics and organisation (Creutz *et al*, 1978). S100 family members are expressed in a cell type-specific manner, often in response to a particular stimulus (Heizmann *et al*, 2002; Marenholz *et al*, 2004). The ANXA2/S100A10 heterotetramer acts as a high affinity F-actin binding protein to regulate the growth of newly formed actin filaments (Hayes *et al*, 2006; Ikebuchi & Waisman, 1990) and thereby controls exo- and endocytosis by actin remodelling (Creutz & Snyder, 2005; Hansen *et al*, 2002). Interestingly, ANXA2 is also identified as a protein induced by ERK5 activation (Fig. 1C), suggesting that ERK5 may regulate the abundance of ANXA2/S100A10 complexes. Our analysis suggests that ERK5 specifically impacts on ANXA2 and S100A10 relative to other family members, which are not significantly altered by ERK5 signalling in our quantitative proteomic analysis (Fig. 1D). Furthermore, mESC protein copy number estimation by absolute quantitative proteomics (Wisniewski *et al*, 2014) indicates that S100A10 (Fig. 1E) and ANXA2 (Fig. 1F) are amongst the most abundant Annexin and S100 family proteins in mESCs, implying that ERK5 signalling may have a functional impact by specifically modulating ANXA2-S100A10 abundance.

### The ERK5 pathway promotes expression of genes encoding S100A10 and ANXA2

As a functional connection between ERK5 signalling and ANXA2-S100A10 has not been reportedly previously, we first set out to validate the role of ERK5 in regulating S100A10 and ANXA2 protein abundance. Analysis of S100A10 and ANXA2 by immunoblotting confirms that ERK5 activation by MEK5DD does indeed lead to augmented S100A10 and ANXA2 protein abundance, and this is reversed by treatment with the ERK5 inhibitor AX15836 (Fig. 2A). This broadly mirrors induction of a known ERK5 transcriptional target, the KLF2 transcription factor (Fig. 2A). We also employed *Erk5/Mapk7*^-/-^ mESCs (Williams *et al*., 2016) to confirm the role of ERK5 in regulation of ANXA2 and S100A10 protein abundance. *Erk5*^-/-^ mESCs transfected with MEK5DD have a basal level of ANXA2 and S100A10 protein (Fig. 2B), which is significantly augmented by re-expression of wild-type (WT) ERK5, but not by a kinase inactive mutant (D200A KD ERK5; Fig. 2B). Thus, these data confirm that S100A10 and ANXA2 are novel targets of the ERK5 signalling pathway in mESCs.

**Figure 2:**
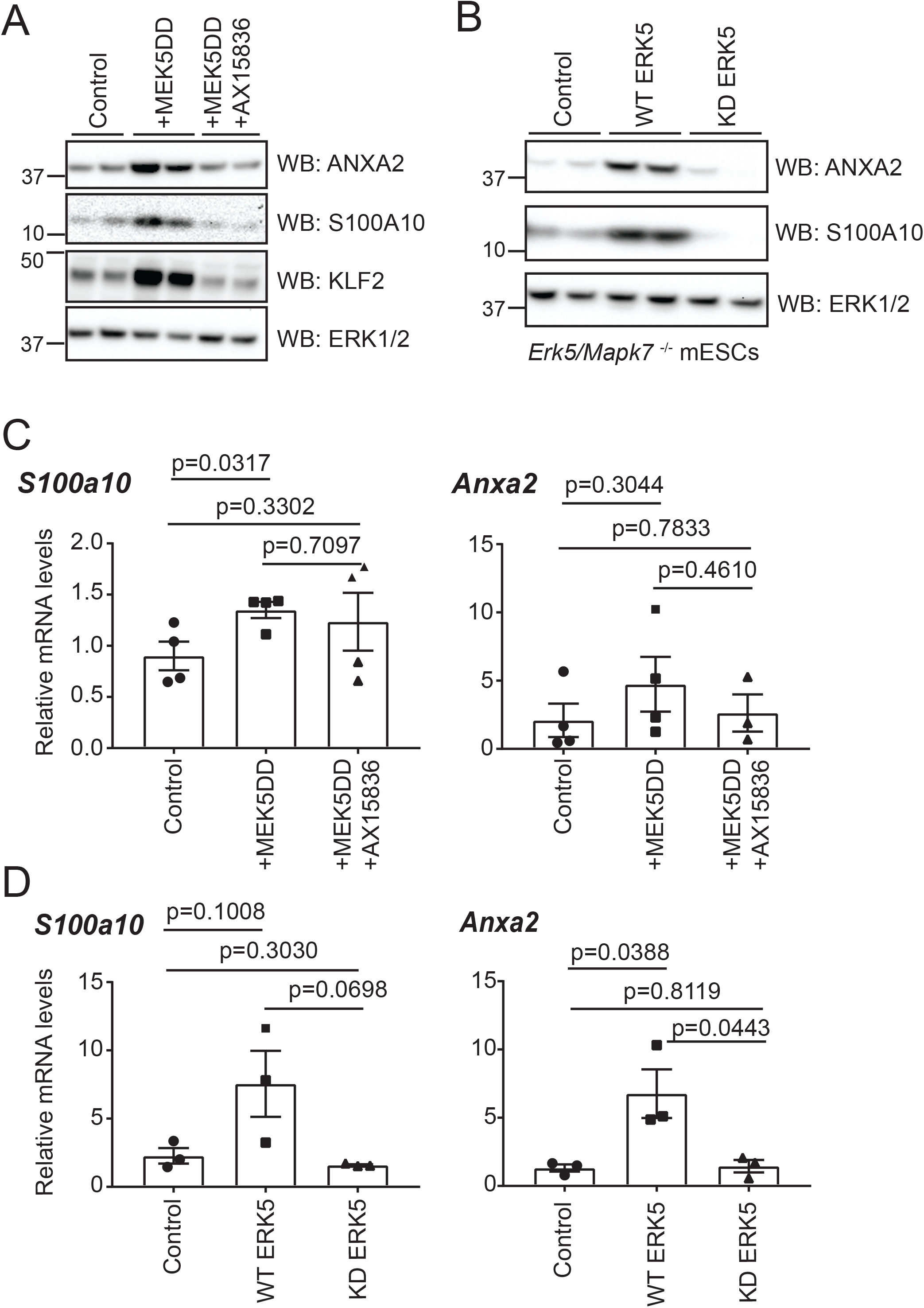
ERK5 signalling regulates S100A10 and ANXA2 protein and gene expression. (A) Wild-type mESCs were transfected with either empty vector (Control), MEK5DD alone (+MEK5DD) to activate the ERK5 pathway, or MEK5DD followed by ERK5 inhibition (+MEK5DD +AX15836). ERK5 inhibition by AX15836 was for 24 h at a final concentration of 10 μM. Cells were lysed and ANXA2, S100A10, and KLF2 levels determined by immunoblotting. ERK1/2 was used as a loading control. Results are representative of 3 independent experiments. (B) *Mapk7* ^-/-^ (ERK5 knock-out) mESCs were transfected with MEK5DD and either empty vector (control), wildtype (WT) ERK5 or a kinase dead (KD) ERK5 mutant (D200A). mESCs were lysed and ANXA2 and S100A10 levels determined by immunoblotting. ERK1/2 was used as a loading control. Results are representative of 3 independent experiments. (C) Wild-type mESCs were transfected with either empty vector (Control), MEK5DD alone (+MEK5DD) to activate the ERK5 pathway, or MEK5DD followed by ERK5 inhibition (+MEK5DD +AX15836). ERK5 inhibition by AX15836 was for 24 h at a final concentration of 10 μM. RNA was extracted and subjected to qRT-PCR analysis. Error bars indicate SEM (n=4). Statistical significance calculated by student’s t-test. (D) *Mapk7* ^-/-^ (ERK5 knock-out) mESCs were transfected with MEK5DD and either empty vector (control), wildtype (WT) ERK5 or a kinase dead (KD) ERK5 mutant (D200A). RNA was extracted and subjected to qRT-PCR analysis. Error bars indicate SEM (n=3). Statistical significance calculated by student’s t-test.

We then sought to determine the mechanism by which ERK5 signalling increases S100A10/ANXA2 protein abundance. As ERK5 plays a key role in transcriptional regulation of pluripotency genes (Williams *et al*., 2016), we hypothesised that the ERK5 pathway augments S100A10/ANXA2 protein abundance by driving expression of the *S100a10* and *Anxa2* genes. Indeed, ERK5 activation by MEK5DD increases *S100a10* and *Anxa2* mRNA levels (Fig. 2C), suggesting that ERK5 signalling promotes *S100a10* and *Anxa2* gene transcription. Furthermore, *S100a10* and *Anxa2* mRNA levels are elevated in *Mapk7/Erk5*^-/-^ mESCs expressing WT ERK5, but not kinase-inactive KD ERK5 (Fig. 2D). Taken together, our data suggest that ERK5 signalling plays a key role in regulating *S100a10* and *Anxa2* gene expression, resulting in increased S100A10/ANXA2 protein abundance in mESCs.

### The ERK5-KLF2 transcriptional axis controls *S100a10* and *Anxa2* gene expression

A key question arising from our results concerns the mechanism by which ERK5 signalling drives *S100A10/ANXA2* gene expression. As shown previously, a major transcriptional target of ERK5 in mESCs is the KLF2 transcription factor (Brown *et al*., 2021; Morikawa *et al*., 2016; Parmar *et al*, 2006; Sunadome *et al*, 2011). Therefore, we hypothesised that transcriptional induction of KLF2 is the mechanism by which ERK5 signalling drives *Anxa2* and *S100a10* transcription. To test this, we used CRISPR/Cas9 gene editing to generate mESCs in which endogenous KLF2 is transiently suppressed (*Klf2*^Δ/Δ^ mESCs; (Brown *et al*., 2021)). In these cells, *S100a10* and *Anxa2* mRNA abundance is robustly enhanced by expression of wild-type KLF2 (Fig. 3A), suggesting that ERK5 signalling regulates *S100a10* and *Anxa2* expression via KLF2-dependent transcriptional induction. Consistent with this notion, ANXA2 and S100A10 protein abundance is also robustly augmented by expression of WT KLF2 (Fig. 3B), and this is further enhanced by expression of a non-phosphorylatable KLF2 mutant that is partially resistant to ubiquitin-mediated proteasomal degradation (4A-KLF2; Fig. 3B) (Brown *et al*., 2021). Finally, elevated ANXA2 and S100A10 protein abundance observed in response to ERK5 activation by MEK5DD is blunted in KLF2-deficient (*Klf2*^Δ/Δ^) mESCs, when compared to wild-type mESCs that express endogenous KLF2 (Fig. 3C). Taken together, our results indicate that the ERK5-KLF2 transcriptional axis is primarily responsible for driving *Anxa2* and *S100a10* gene expression, leading to increased abundance of ANXA2-S100A10 in mESCs.

**Figure 3:**
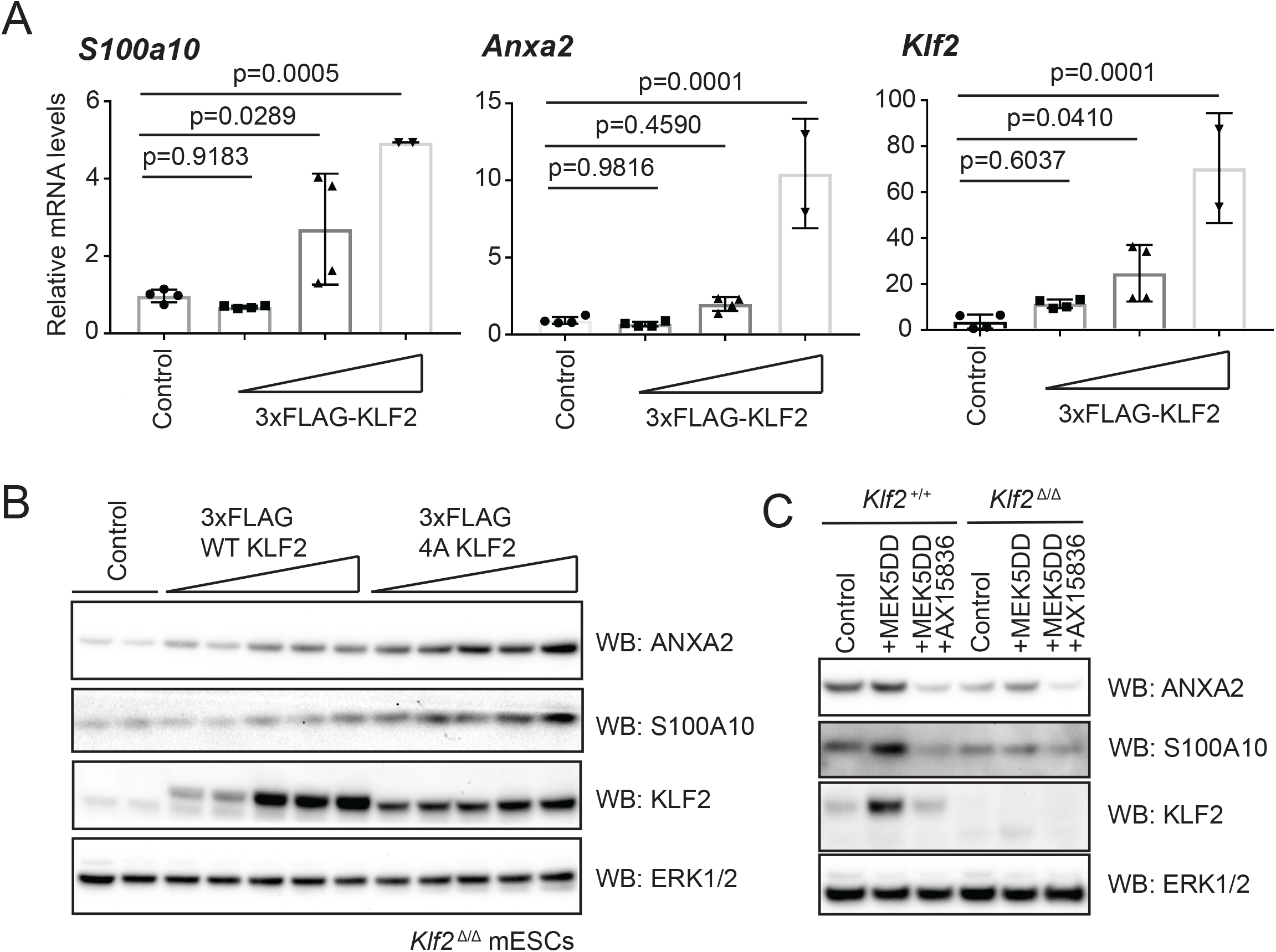
ERK5 signalling regulates S100A10 and ANNEXIN A2 protein and gene expression. (A) *Klf2*^Δ/Δ^ mESCs were transfected with either empty vector (Control) or increasing concentrations of 3xFLAG-KLF2. mESCs were lysed, RNA extracted and subjected to qRT-PCR analysis. Error bars indicate SEM (n=4, n=2 for the highest amount of 3xFLAG KLF2 cDNA [0.5μg]). Statistical significance calculated by one-way ANOVA with multiple comparisons. (B) *Klf2*^Δ/Δ^ mESCs were transfected with either empty vector (Control) or increasing concentrations of 3xFLAG-KLF2 wild-type (WT) or a 4x Ser-Ala mutant (4A). mESCs were lysed and ANXA2, S100A10 and KLF2 levels were determined by immunoblotting. ERK1/2 was used as a loading control. Results are representative of 3 independent experiments. (C) *Klf2*^+/+^ and *Klf2*^Δ/Δ^ mESCs were transfected with either empty vector (Control), MEK5DD alone (+MEK5DD) to activate the ERK5 pathway, or MEK5DD followed by ERK5 inhibition (+MEK5DD +AX15836). ERK5 inhibition by AX15836 was for 24 h at a final concentration of 10 μM. mESCs were lysed and ANXA2, S100A10, and KLF2 levels determined by immunoblotting. ERK1/2 was used as a loading control. Results are representative of 3 independent experiments.

### KLF transcription factors bind to *S100a10* and *Anxa2* gene regulatory elements

In order to address the mechanism and specificity of *Anxa2* and *S100a10* gene regulation by KLF2, we next investigated whether KLF2 and other KLF family members engage regulatory elements of S100 and ANXA family genes. To this end, we performed mining of chromatin immunoprecipitation followed by DNA sequencing (ChIP-SEQ) data stored in the CODEX next generation sequencing database (Sanchez-Castillo *et al*, 2015). This analysis revealed that KLFs bind to multiple sites in close proximity to transcriptional start sites of *S100* (Fig. 4A) and *ANXA* (Fig. 4B) family genes, suggesting that they could be targets for transcriptional regulation. Interestingly, although KLF2 engages the several S100 and ANXA encoding genes, multiple KLF2 binding sites are found proximal to the *S100a10* and *Anxa2* transcriptional start sites (Fig. 4A,B), consistent with the notion that KLF2 is an important regulator of their expression.

**Figure 4:**
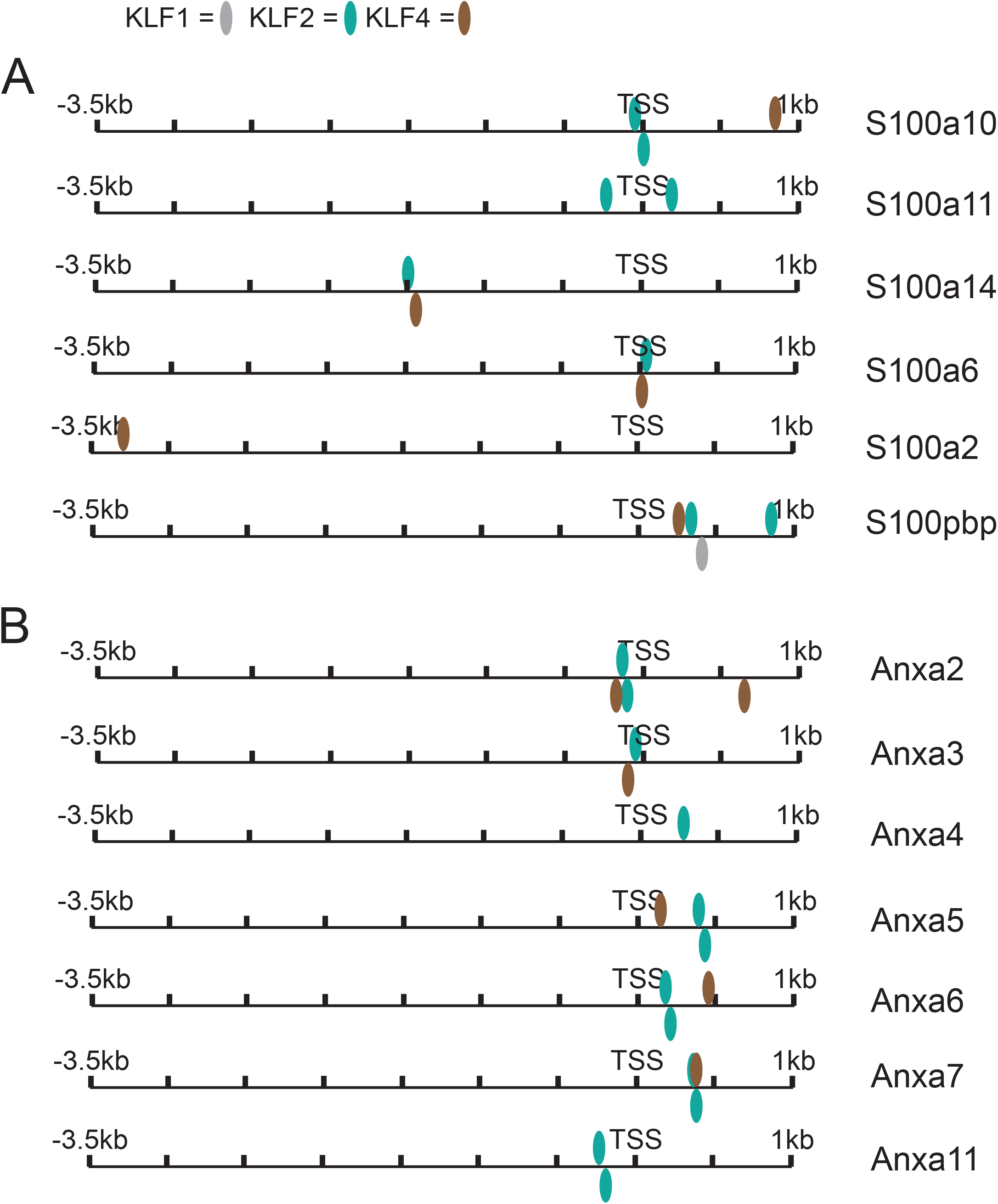
KLF family transcription factors bind proximal to Annexin and S100 promoters. (A) Data extracted from CODEX database (Sanchez-Castillo *et al*, 2015) for regions immediately 5’ and 3’ of the transcriptional start site (−3.5 kb to +1.0 kb) of S100 family genes. The indicated S100 and KLF family members are those represented in the CODEX database. (B) Data extracted from CODEX database for regions immediately 5’ and 3’ of the transcriptional start site (−3.5 kb to +1.0 kb) of Annexin family genes. The indicated Annexin and KLF family members are those represented in the CODEX database.

### The ERK5 pathway drives ANXA2-S100A10 complex assembly

As ERK5 signalling promotes *S100a10* and *Anxa2* gene expression leading to increased protein abundance, we hypothesised that ERK5 will promote formation of intact S100A10-ANXA2 complexes. Co-immunoprecipitation of endogenous ANXA2 and S100A10 proved challenging, possibly due to antibody epitopes mapping to the S100A10-ANXA2 interface. To circumvent this, we expressed N-terminally tagged constructs of ANXA2 (HA) and S100A10 (3xFLAG) and performed FLAG immunoprecipitation to pull down S100A10 complexes. As expected, 3xFLAG-S100A10 is pulled down in both the presence and absence of overexpressed HA-ANXA2 (Fig. 5A), although less 3xFLAG-S100A10 is expressed in the absence of overexpressed HA-ANXA2, consistent with a report that ANXA2 complexing protects S100A10 from ubiquitin-mediated degradation (He *et al*, 2008). Furthermore, ANXA2 (HA-tagged or endogenous) complexing with 3xFLAG-S100A10 is detected only at higher levels of 3xFLAG-S100A10 and HA-ANXA2 expression levels (Fig. 5A, compare lanes 3 and 4). This data suggests that S100A10-ANXA2 complex formation is observed only at high levels of S100A10 and ANXA2 expression, which in mESCs is achieved by activation of the ERK5-KLF2 signalling axis.

**Figure 5:**
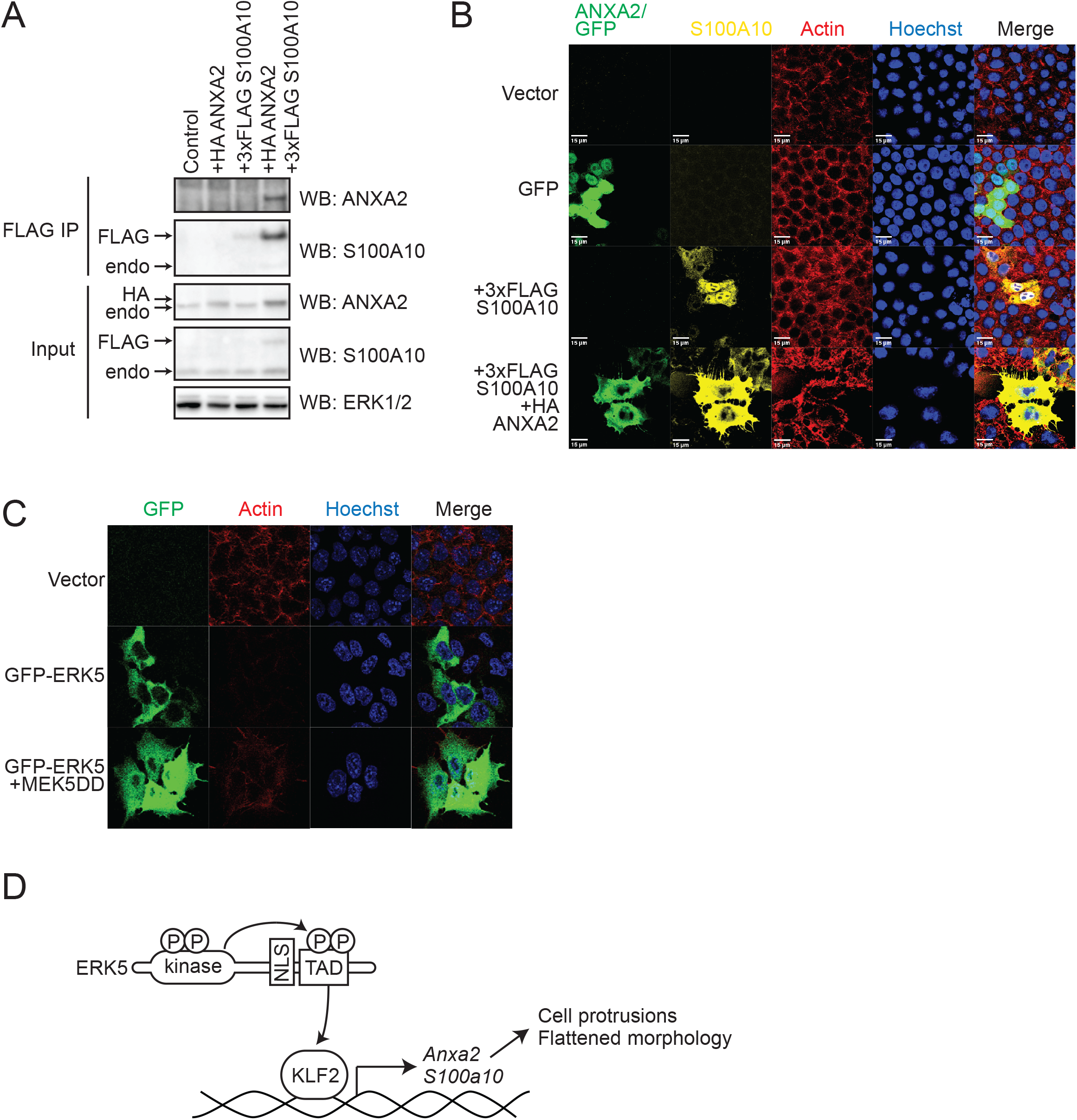
Formation of ANXA/S100A10 complex results in a change in cell morphology. (A) Wild-type mESCs were transfected with either empty vector (Control) 3xFLAG-S100A10, HA-ANXA2 or both, lysed and FLAG IP was used to isolate S100A10 complexes. ANXA2 and S100A10 levels were determined by immunoblotting. ERK1/2 was used as a loading control for input samples. (B) Wild-type mESCs were transfected with either empty vector (Control), GFP, 3xFLAG-S100A10, or 3xFLAG-S100A10 with HA-ANXA2 for 24 hr with no puromycin selection prior to fixing with 4% PFA. GFP, ANXA2, and S100A10 were visualised by immunofluorescence. ActinRed stain was used to show actin cytoskeleton, and Hoechst stain was used to show nuclei. GFP control was used to visualise whether mESC transfection results in a morphology change that is unrelated to ANXA2/S100A10 expression. Scale bar = 15μm (C) mESCs were transfected with either empty vector (Control), GFP-ERK5 or GFP-ERK5 +MEK5DD prior to fixing in 4% PFA. GFP-ERK5 was visualised by immunofluorescence, ActinRed stain was used to show actin cytoskeleton, and Hoechst stain was used to show nuclei. (D) Model for ERK5-depdendent regulation of ANXA2/S100A10. ERK5 activation and transcriptional induction of KLF2 leads to increased *Anxa2/S100a10* gene expression, which drives cellular morphology characterised by flattening and protrusions.

### ERK5 signalling and ANXA2-S100A10 complex assembly drives mESC membrane dynamics

ANXA2 and S100A10 have been implicated in regulation of cellular trafficking and membrane dynamics. Therefore, we investigated the impact of elevated ANXA2-S100A10 abundance in response to ERK5 signalling on mESC morphology. mESCs expressing either control vector or a vector expressing GFP exhibit a tightly packed colony morphology, as visualised by actin (ActinRed) and DNA staining (Hoechst; Fig. 5B). This morphology is also observed upon expression of 3xFLAG-S100A10 in mESCs (Fig. 5B). However, co-expression of 3xFLAG-S100A10 with HA-ANXA2 leads to flattened mESC morphology accompanied by formation of large membrane protrusions reminiscent of filopodia (Fig. 5B). These data suggest that elevated expression of ANXA2-S100A10 complexes in response to ERK5 signalling may lead to altered mESC morphology characterised by flattening and formation of membrane protrusions. Importantly, this phenotype is characteristic of mESCs in which ERK5 signalling is activated, as mESCs expressing either control vector or a vector expressing GFP-ERK5 alone exhibit a tightly packed colony morphology (Fig. 5C), whilst mESCs expressing GFP-ERK5 activated by MEK5DD exhibit a similar, flattened morphology with membrane protrusions (Fig. 5C). Taken together, our results suggest that ERK5-dependent induction of ANXA2-S100A10 complexes can drive altered mESC morphology.

## Discussion

ERK5 controls expression of pluripotency genes and stem cell rejuvenation factors in mESCs (Brown *et al*., 2021; Morikawa *et al*., 2016; Williams *et al*., 2016). However, further transcriptional targets of the ERK5 pathway in pluripotent cells have not been identified. Here, we employ quantitative proteomics to identify ANXA2-S100A10 as major transcriptional targets of the ERK5 pathway. We show that ERK5 signalling specifically increases the abundance of ANXA2 and S100A10 proteins, while other Annexin and S100 family members are largely unaffected. We identify KLF2 as a key transcription factor that promotes ANXA2-S100A10 expression, likely via direct transcriptional regulation (Fig. 5D). We also show that ERK5 signalling and ANXA2-S100A10 induction lead to a flattened cell morphology and altered membrane dynamics characterised by formation of membrane protrusions, which we hypothesise are filopodia. These findings uncover a potential function for ERK5 and ANXA2-S100A10 in regulating cellular morphology and structure. Future work will examine the identity of these structures and explore specific impacts of ERK5 signalling on cell migration and motility.

These findings also suggest a novel function of ERK5 that is relevant for development of cancer. Previous data implicate ANXA2-S100A10-dependent membrane dynamics in regulation of key physiological and pathophysiological processes such as motility, invasion, metastasis, and epithelial to mesenchymal transition (EMT). As such, the cell morphology changes observed upon ERK5 induction of ANXA2-S100A10 prompts the hypothesis that ERK5 may play a role in their regulation. Indeed, ERK5 is known to drive aggressive cancers with high rates of metastasis (Pavan *et al*, 2018; Ramsay *et al*, 2011), a phenotype also associated with expression of ANXA2-S100A10 in tumours (Bharadwaj *et al*, 2013; Heizmann *et al*., 2002). Future studies will investigate the importance of ERK5 regulation of ANXA2-S100A10 in various relevant developmental and disease contexts.

## Materials & Methods

Many reagents developed for this study are available by request at the MRC-PPU reagents & services website (https://mrcppureagents.dundee.ac.uk/).

### Mass spectrometry analysis

Quantitative total proteomic mass spectrometry analysis was reported previously and detailed methods provided in (Brown *et al*., 2021). Briefly, protein extraction and tryptic digestion was followed by TMT isobaric labelling, high pH reverse phase fractionation and LC–MS/MS analysis.

### Antibodies and chemicals

All antibodies used in this study are listed in Table 1 below. FLAG-M2 agarose (Sigma) was used for FLAG immunoprecipitation. AX15836 (Tocris) was used as an ERK5 inhibitor.

**Table 1.** Antibodies.

| TARGET | SOURCE | CAT. NO. | DILUTION | HOST |
| --- | --- | --- | --- | --- |
| KLF2 | Merck Millipore | 09-820 | 1:1000 | Rabbit |
| ERK1 | Santa Cruz Biotechnology | Sc-93 | 1:1000 | Rabbit |
| ANNEXIN A2 | BD | 610068 | 1:1000 | Mouse |
| S100A10 | R&D Systems | AF2377 | 1:1000 | Goat |
| FLAG (HRP) | Sigma | A8592-2 | 1:1000 |  |
| HA (HRP) | Roche | 12013819001 | 1:1000 |  |
| GFP | Abcam | Ab13970 | 1:500 | Chicken |

### cDNA constructs

All cDNA constructs used in this study are listed in Table 2 below and are available by request at the MRC-PPU reagents & services website (https://mrcppureagents.dundee.ac.uk/).

**Table 2.** cDNA constructs.

| CDNA | TAG | VECTOR | CODE | RESISTANCE MARKER |
| --- | --- | --- | --- | --- |
| ERK5 | No tag | pCAGGS | DU50514 | Puromycin |
| ERK5 D200A | No tag | pCAGGS | DU50662 | Puromycin |
| GFP-ERK5 | N-terminal GFP | pCAGGS | DU50830 | Puromycin |
| **MEK5 S311D T315D** | N-terminal 3xFLAG | pCAGGS | DU53252 | Puromycin |
| **KLF2** | N-terminal 3xFLAG | pCAGGS | DU56075 | Puromycin |
| **KLF2 T171A S175A S243A T247A (4A)** | N-terminal 3xFLAG | pCAGGS | DU53945 | Puromycin |
| **ANXA2** | N-terminal HA | pCAGGS | DU61146 | Puromycin |
| **ANXA2** | C-terminal HA | pCAGGS | DU61136 | Puromycin |
| **S100A10** | N-terminal FLAG | pCAGGS | DU61138 | Puromycin |
| **S100A10** | C-terminal FLAG | pCAGGS | DU61134 | Puromycin |
| **GFP** |  | pCAGGS |  | Puromycin |

### mESC culture, transfection and lysis

CCE mESCs were cultured on gelatin-coated plates in media containing LIF, 10% FBS (Gibco), and 5% knockout serum replacement (Invitrogen). *Mapk7/Erk5*^-/-^ (Williams *et al*., 2016) and *Klf2*^Δ/Δ^ (Brown *et al*., 2021) mESCs were reported previously. Typically, mESCs were transfected with pCAGGS cDNA expression vectors using Lipofectamine LTX (Life Technologies) and 24 h later selected with puromycin for a further 24 h. Cell extracts were made in lysis buffer (20 mM Tris [pH 7.4], 150 mM NaCl, 1 mM EDTA, 1% NP-40 [v/v], 0.5% sodium deoxycholate [w/v], 10 mM β-glycerophosphate, 10 mM sodium pyrophosphate, 1 mM NaF, 2 mM Na_3_VO_4_, and Roche Complete Protease Inhibitor Cocktail Tablets). Immunoprecipitation

### qRT-PCR

RNA was extracted using the OMEGA total RNA kit and reverse transcribed using iScript reverse transcriptase (Bio-Rad). qPCR was performed using TB Green Premix Ex Taq (Takara). The ΔCt method using *Gapdh* as a reference gene was used to analyse relative expression and the 2^−ΔΔCt^ (Livak) method used to normalise to control. Primers used are listed in Table 3 below.

**Table 3.**
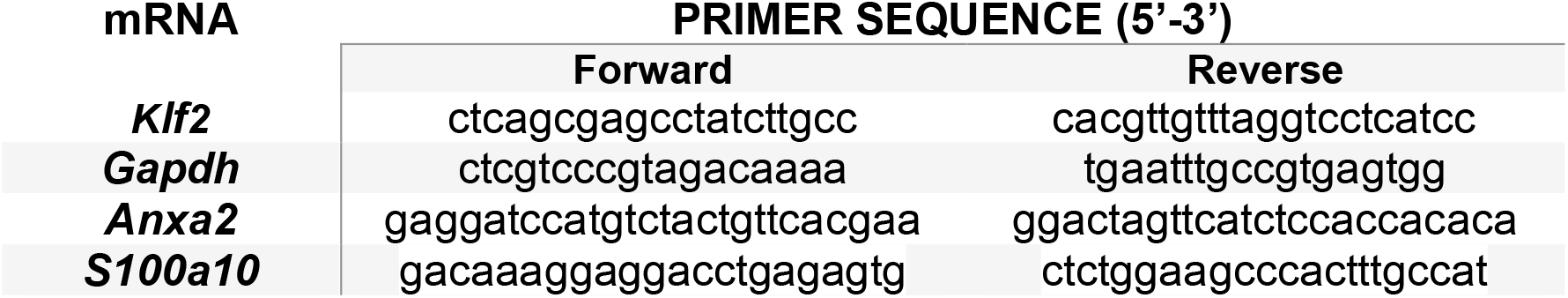
qRT-PCR primer sequences.

### CODEX

Chromatin immunoprecipitation followed by DNA sequencing data for genomic binding sites of KLF family transcription were extracted from the CODEX database (Sanchez-Castillo *et al*, 2015) for regions immediately 5’ and 3’ of the transcriptional start site (−3.5 kb to +1.0 kb) of Annexin and S100 family genes. The indicated Annexin, S100 and KLF family members are those reported in the CODEX database.

### Immunofluorescence

mESCs were seeded on gelatin-coated coverslips and fixed with 4% PFA (w/v) in PBS, before being permeabilised in 0.5% Triton X-100 (v/v) in PBS for 5 mins at room temperature. Coverslips were then blocked with 1% fish gelatin (w/v) in PBS and incubated with primary antibody for 2 h at room temperature. After three washes with PBS, fluorescent secondary antibodies were diluted 1:500 in blocking buffer and incubated on coverslips for 1 h at room temperature. Actin Red 555 reagent was added to the secondary antibody mix. After three washes with PBS, 0.1 µg/ml Hoechst was incubated with the coverslips for 5 min at room temperature in a humid chamber to stain nuclei. After three more washes with PBS, coverslips were mounted onto cover slides using Fluorsave reagent (Millipore). Images were taken using a Leica SP8 confocal microscope and processed using FIJI and Photoshop CSC software (Adobe).

## Data Availability

Mass spectrometry proteomics data have been deposited to the ProteomeXchange Consortium via the EBI PRIDE repository with the dataset identifier PXD024679. Other data generated and/or analysed during the current study are available from the corresponding authors on request.

